# Structural analysis of 1-deoxy-D-xylulose 5-phosphate synthase from *Pseudomonas aeruginosa* and *Klebsiella pneumoniae* reveals conformational changes upon cofactor binding

**DOI:** 10.1101/2022.07.04.498669

**Authors:** Rawia Hamid, Sebastian Adam, Antoine Lacour, Leticia Monjas Gomez, Anna K. H. Hirsch

## Abstract

Isoprenoid precursor biosynthesis is an essential part of primary metabolism in all living organisms. While eukaryotes utilize the mevalonate (MEV) pathway for isoprenoid-precursor biosynthesis, the most important bacterial pathogens rely on the methylerythritol-phosphate (MEP) pathway. Therefore, enzymes involved in the MEP pathway are potentially valuable targets for the development of novel antibacterials to tackle the current antimicrobial resistance crisis. Within the MEP pathway, the enzyme 1-deoxy-D-xylulose-5-phosphate synthase (DXPS) represents a crucial, rate-limiting first step and a branch point in the biosynthesis of the vitamins B1 and B6. Herein, we present two novel, high-resolution DXPS crystal structures of the important ESKAPE pathogens *Pseudomonas aeruginosa* and *Klebsiella pneumoniae* in both the co-factor-bound and apo forms. We demonstrate that the absence of the cofactor ThDP leads to a disordered loop close to the active site and may be important for the design of potent DXPS inhibitors, albeit being different in both structures. In addition, we report the complex structure of paDXPS with a fluoropyruvate adduct, shedding more light on the structural basis of DXPS catalysis. Lastly, we have determined a complex structure of paDXPS with a thiamine analogue, opening up a route for structure-based drug design of this essential enzyme of the MEP pathway.

## Introduction

The widespread use of antibiotics in the past five decades has caused bacteria to develop resistance mechanisms to evade the detrimental effect of these agents ^1^. This effect is especially critical for infections caused by ESKAPE pathogens, which are difficult to treat and at times even fatal. *Pseudomonas aeruginosa and Klebsiella pneumoniae* are both Gram-negative pathogens with the ability to integrate exogenous DNA to obtain antibiotic resistance (horizontal gene transfer)^2^. Both organisms can adapt to the environment of human airways^3^ and are major causes of opportunistic infections, predominantly, pneumonia and sepsis in hospitalized patients and patients with cystic fibrosis^4,5^. Measures such as antibiotic stewardship programs to improve the way antibiotics are prescribed and synthesis of novel chemical entities with new modes of action need to be implemented in order to tackle the antimicrobial-resistance crisis.

The methylerythritol-phosphate (MEP) pathway has attracted attention as it represents a rich source of potentially attractive anti-infective targets. The MEP pathway, also called the mevalonate-independent pathway, is considered to be the main pathway for the synthesis of isoprenoid building blocks in most bacteria, plants and protozoa ^6–8^. Its absence in animals makes the enzymes of this pathway particularly interesting targets for drug discovery. The complete pathway consists of seven enzymes, each of which could be targeted individually. 1-Deoxy-D-xylulose-5-phosphate synthase (DXPS) catalyzes the first, rate-limiting step of the MEP pathway. The enzyme is responsible for the formation of 1-deoxy-D-xylulose 5-phosphate (DXP) by condensation of pyruvate and glyceraldehyde 3-phosphate (GAP) in the presence of thiamine diphosphate (ThDP) and magnesium as co-factors. DXP is an important metabolite in not only the biosynthesis of isoprenoid precursors, but also in vitamin B1 (thiamine) and vitamin B6 (pyridoxine) biosynthesis^9^.

The first DXPS structures of *Deinococcus radiodurans* and *Escherichia coli* were solved via X-ray crystallography and published in 2007 ^10^. Subsequent studies on these structures have shown that DXPS undergoes conformational changes during catalysis ^11–13^. Meyers *et al*. devised the name spoon-fork motif to describe a loop movement involved in binding of a post-decarboxylation intermediate formed by interaction of pyruvate with the ThDP-bound enzyme followed by D/L-GAP^11^. Thus far, few experimentally determined structures of DXPS have been reported^14^, but due to the generally conserved active sites, the available crystal structures of DXPS are used as a model for structure-based drug design^12,15,16^. However, this has proven to be very limiting, as three-dimensional active sites can typically not be deduced from sequence homology alone. We hereby report the first crystal structures of both the apo and cofactor-bound forms of *P. aeruginosa* and *K. pneumoniae* DXPS. We demonstrate that binding of ThDP leads to conformational changes involving a loop in domain I that is involved in the shaping of the active site, which also contributes to the binding of diphosphate. In addition, we have determined the DXPS structure of *P. aeruginosa* with an intermediate formed by the interaction of paDXPS and the substrate analogue fluoropyruvate. Lastly, we have determined the crystal structure of paDXPS with a thiamine analogue. We therefore provide a structural analysis to support our enzymatic studies and drug-design efforts targeting this important enzyme from the MEP pathway.

## Results and discussion

### Expression, purification and kinetic characterization of paDXPS and kpDXPS

Our group recently developed a truncation strategy to remove a flexible loop in *D. radiodurans* (drDXPS) to produce a protein that crystallizes readily and produces high-quality crystals that diffract to a resolution better than 2.1 Å^17^. Given the success of this construct to produce a more rigid protein and the decent sequence homology between DXPS enzymes from different organisms, we investigated the applicability of this strategy to other homologues. As a result, we designed a mutated version of paDXPS and kpDXPS *in silico*, in which we replaced a flexible loop with six glycine units. This loop typically has a very low evolutionary conservation and does neither shape the active site nor is it involved in the formation of secondary structure^17^. After multiple sequence alignments of published sequences of paDXPS and kpDXPS, the loop was identified to be corresponding to residues 206–245 in paDXPS and residues 197–240 in kpDXPS. The respective genes for paDXPS and kpDXPS were obtained from BioCat and transformed into *E. coli* BL21 (DE3) or other BL21 derivatives. The enzymes were heterologously expressed, purified and analyzed concerning their purity and identity. Next, we analyzed the effect of this mutation on the activity of new homologue constructs and evaluated and compared the enzyme kinetics for both versions of the enzyme. The results summarized in Table 1 demonstrate comparable activities of the native (paDXPS) and truncated (ΔpaDXPS) enzymes, with a tendency for ΔpaDXPS to show slightly higher affinities for both substrates pyruvate and D/L-GAP. Although the native kpDXPS was not tested, the mutated version is showing reasonable kinetics for both substrates.

**Table 1:** The steady-state kinetic parameters of pyruvate and DL-GAP for the wild-type and truncated enzymes paDXPS and mutated kpDXPS.

| Enzyme | $K_m$ D-GAP (μM) | $K_m$ Pyruvate (μM) |
| --- | --- | --- |
| paDXPS | $312 \pm 104$ | $133 \pm 8.7$ |
| ΔpaDXPS | $228 \pm 98$ | $80 \pm 5.6$ |
| ΔkpDXPS | $149.7 \pm 6.3$ | $77 \pm 16.1$ |

After confirming the identity and activity of our purified enzymes, we opted to elucidate the structures of these novel homologues from ESKAPE pathogens.

### Overall structural analysis of paDXPS from *Pseudomonas aeruginosa*

DXPS from *Pseudomonas aeruginosa* was purified as described in the methods section and crystallization trials were carried out with commercially available screens. Crystals grew after 3–5 days in a number of conditions, after which several optimization trials were performed. Crystals were obtained at 18 °C using the hanging drop vapor-diffusion method with a reservoir solution consisting of 100 mM HEPES, 12% PEG 8000 and 200 mM calcium acetate and a ΔpaDXPS concentration of 15 mg/mL. Single crystals were mounted, and datasets were collected at the Swiss Light Source up to a resolution limit of 2.0Å. The structure was initially determined with molecular replacement using the published structure of drDXPS (PDB ID: 6OUV) as a search model. After that, a refined paDXPS structure was used as the search model for subsequent structures. All data collection and refinement statistics can be found in the supplementary Table 2, and the structures were deposited in the protein data bank. In total, we report the structures of ThDP-bound paDXPS, an apo structure, a complex structure with the substrate analogue fluoropyruvate and a complex structure with a selected ThDP analogue in subsequent paragraphs.

In general, *pa*DXPS crystallized in orthorhombic crystal form (space group P212121) and diffracted up to a resolution limit of 2.0 Å. The asymmetric unit consists of six protein chains with a distribution into two dimeric units and two monomeric units. In most obtained paDXPS structures, the first 30 N-terminal residues are missing due to a lack of density, which did not affect the correct molecular folding of the complete structure. A native mass spectrometry (MS) measurement performed on paDXPS confirmed that the enzyme exists as a dimer in solution (Fig. S2), which is consistent with the crystallographic results and suggests that the dimer is the biological unit *in vivo*.

PaDXPS shares high sequence homology with homologues from other bacteria, with sequence identities ranging from 43.3% (*D. radiodurans*) and 58.6% (*E. coli*) to 62.5% (*K. pneumoniae*) (Fig. S1). Expectedly, the overall structure of paDXPS is also highly similar to other previously identified homologues from *E. coli* (ecDXPS) and *D. radiodurans* ^10^ (drDXPS), with measured root mean square deviations (RMSD) of 0.785 Å (2881 to 2881 atoms) with drDXPS and 0.643 Å (2914 to 2914 atoms) with ecDXPS (Fig. S1, 2). Each paDXPS monomer consists of three relatively separated domains. Domain I (residues 1–306) consists of five parallel β-sheets and five α-helices, domain II (residues 317–493) contains six parallel β-sheets and six α-helices and domain III (residues 496–620) is built up of five β–sheets, with the first strand being anti-parallel to the other four strands and five α-folds (Fig. 1). The loop connecting domains I and II (residues 307–315) is highly flexible and appears to be disordered in most structures. However, when compared to the structure of drDXPS (PDB ID: 2O1X) (Fig. S1), paDXPS shows some structural differences, e.g., the β-sheet number 5 in domain I is significantly shorter, comprising only three residues (251–253) compared to eight residues in drDXPS. Moreover, the loop spanning residue 307–316, albeit not complete, shows a more open conformation when comparing the ThDP-bound structures.

**Figure 1:**
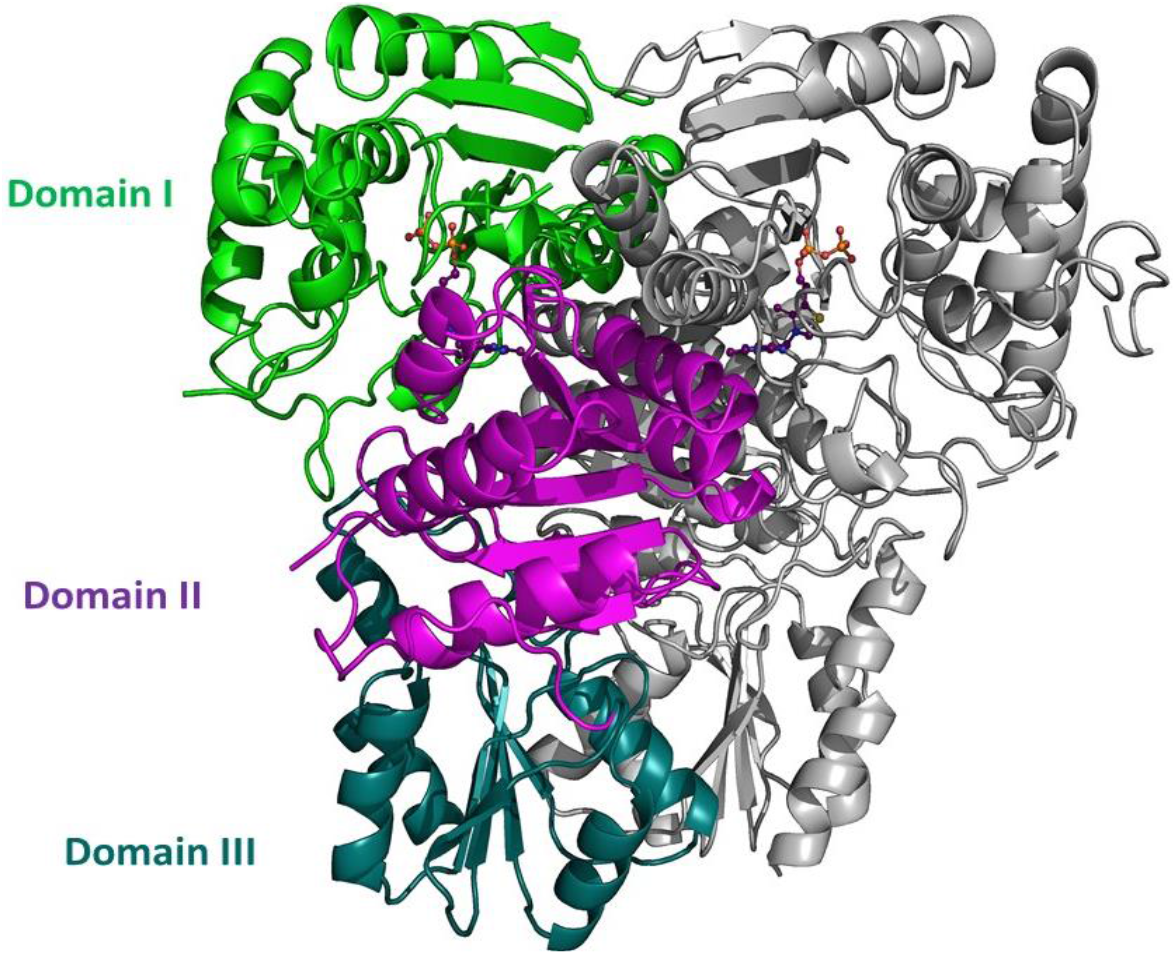
paDXPS-ThDP 3D structure of a dimer showing domain organization (PDB ID: **8A5K**): domain I (green; residues 1–306) contains a five-stranded parallel β-sheet, domain II (violet; residues 317–493) contains a six-stranded parallel β-sheet, domain III (cyan; residues 496–620) contains a five-stranded β-sheet:. All figures of this type were created in PyMol.

### Structural analysis of ThDP-bound paDXPS

Given that the binding site of the cofactor ThDP is the most promising site for drugs targeting ThDP-dependent enzymes including DXPS ^12,15,16^, it is vital to understand the precise location of ThDP and the general three-dimensional shape of the pocket. It is also considered the main druggable pocket in DXPS enzymes^18^, as the folding of the protein leads to an almost complete solvent exposure of the surface, with the exception of the narrow active site. We therefore determined the high-resolution crystal structure of *P. aeruginosa* DXPS in complex with its cofactor ThDP. Co-crystals of ThDP-bound paDXPS were obtained by incubation of 15 mg/mL paDXPS with 500 μM ThDP and 500 μM MgCl for 2h at RT, after which crystals grew in hanging drops with the known condition. A complete dataset was collected at SLS beamline PXIII (X06DA), the structure was solved using molecular replacement and refined to 2.37 Å (PDB ID: 8A5K).

Like other reported homologues ^10^, the binding site of ThDP is located at the interface between domains I and II with the pyrimidine ring facing domain II and the diphosphate moiety facing domain I. In addition, the loop between residues 216–220 in domain I is stabilized by the interaction with the diphosphate and a magnesium cation. ThDP adopts a chair-like conformation; the pyrimidine ring is perpendicular to the thiazole ring, which is, in turn, perpendicular to the diphosphate. The pyrimidine ring of ThDP is π-π stacked against Phe395, while N3 of the pyrimidine is forming a direct backbone interaction with Ser158 (Fig. 2). Interestingly, this residue is replaced by alanine in drDXPS and does not seem to be essential. On the other hand, N1 of the pyrimidine is forming a hydrogen bond with Pro344 and the diphosphate/Mg^2+^ moiety is coordinated by residues Gly188, Gly186, His115, Asn216 and Lys286 (Fig. 4). In total, the paDXPS structure shows ten amino acid mutations in the binding site when compared to the structure of drDXPS (Fig. 3). From these mutations, only amino acids in positions 87, 158 and 189 are forming a direct interaction with ThDP. Of these, the most noteworthy is the mutation of Ala to Ser158, where the hydroxyl group of the serine side chain can possibly be targeted in drug design to form a hydrogen bond with a ligand.

**Figure 2:**
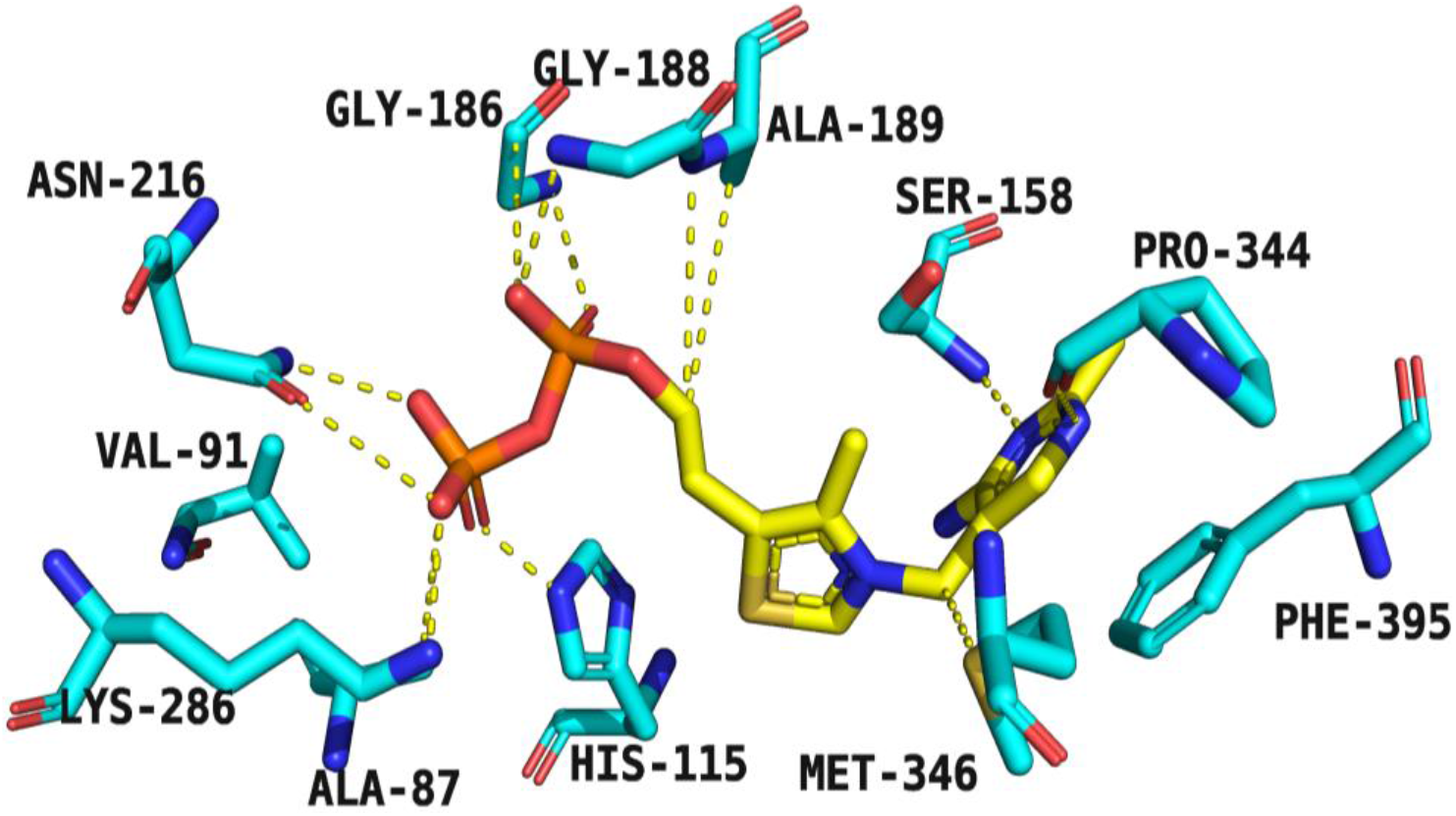
paDXPS active site view showing residues involved in binding to ThDP (PDB ID: **8A5K**). ThDP atoms are shown as sticks: (C, yellow; O, red; N, blue; P, orange), residue atoms are shown as sticks (C, cyan; O, red; N, blue; P, orange; S, yellow). Bonding interactions are shown as yellow dashed lines.

**Figure 3:**
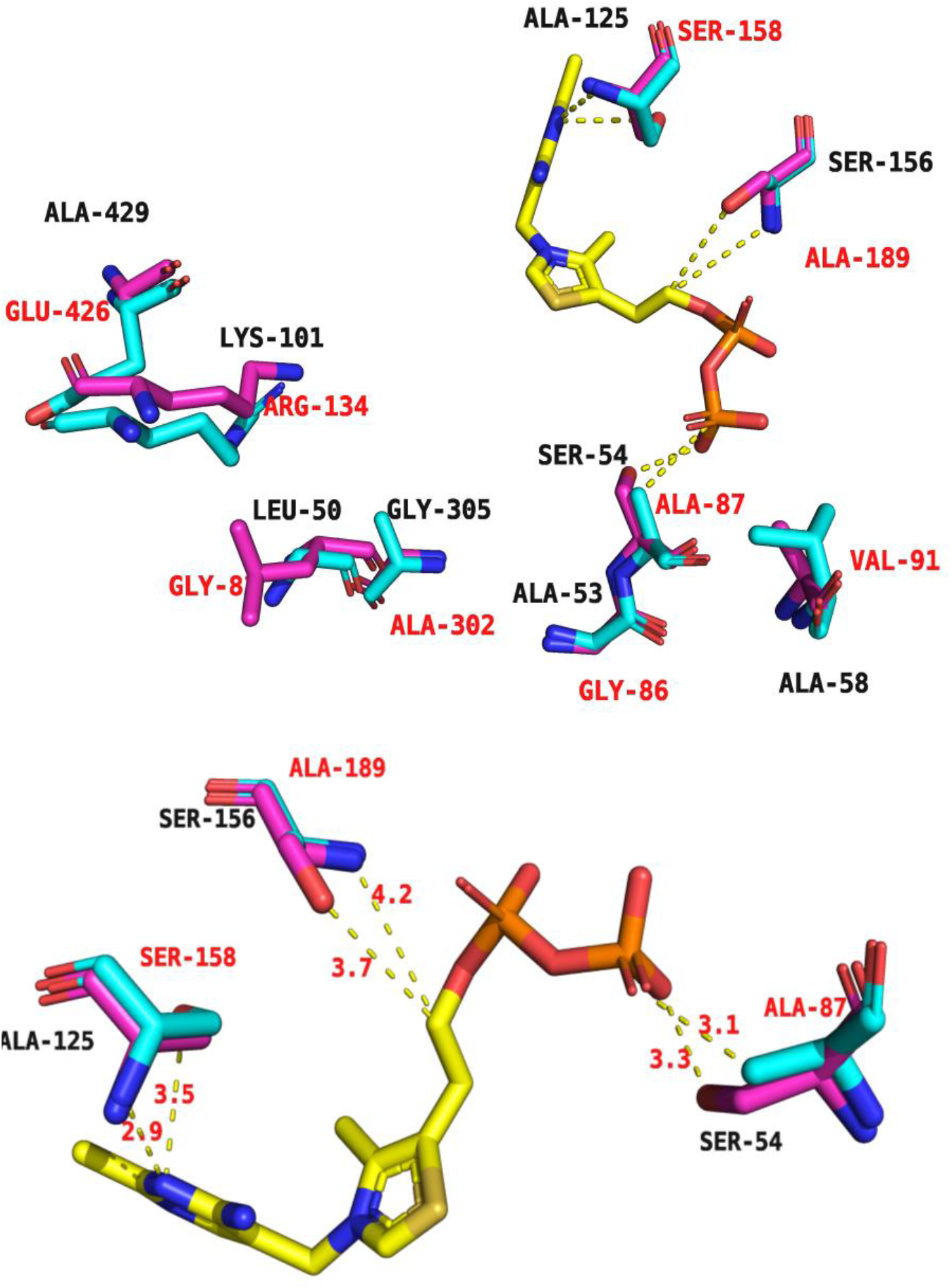
Substrate channel highlighting mutations in/near the active site, paDXPS structure (PDB ID: **8A5K)** is shown in cyan, superimposed with the drDXPS structure in magenta, mutations in paDXPS structure residues are labeled in red: Ala to Ser158, Ser to Ala189, Ala to Ser54,. Mutations near but not included in the active site: Ala to Gly86, Ser to Ala87, Ala to Val91, leu to Gly83Lys to Arg134, Ala to Glu426, Gly to Ala302. ThDP atoms are shown as sticks: (C, yellow; O, red; N, blue; P, orange), protein atoms are shown as sticks (C, cyan; O, red; N, blue; P, orange; S, yellow). Interactions are shown as yellow dashed lines (below 5Å).

**Figure 4:**
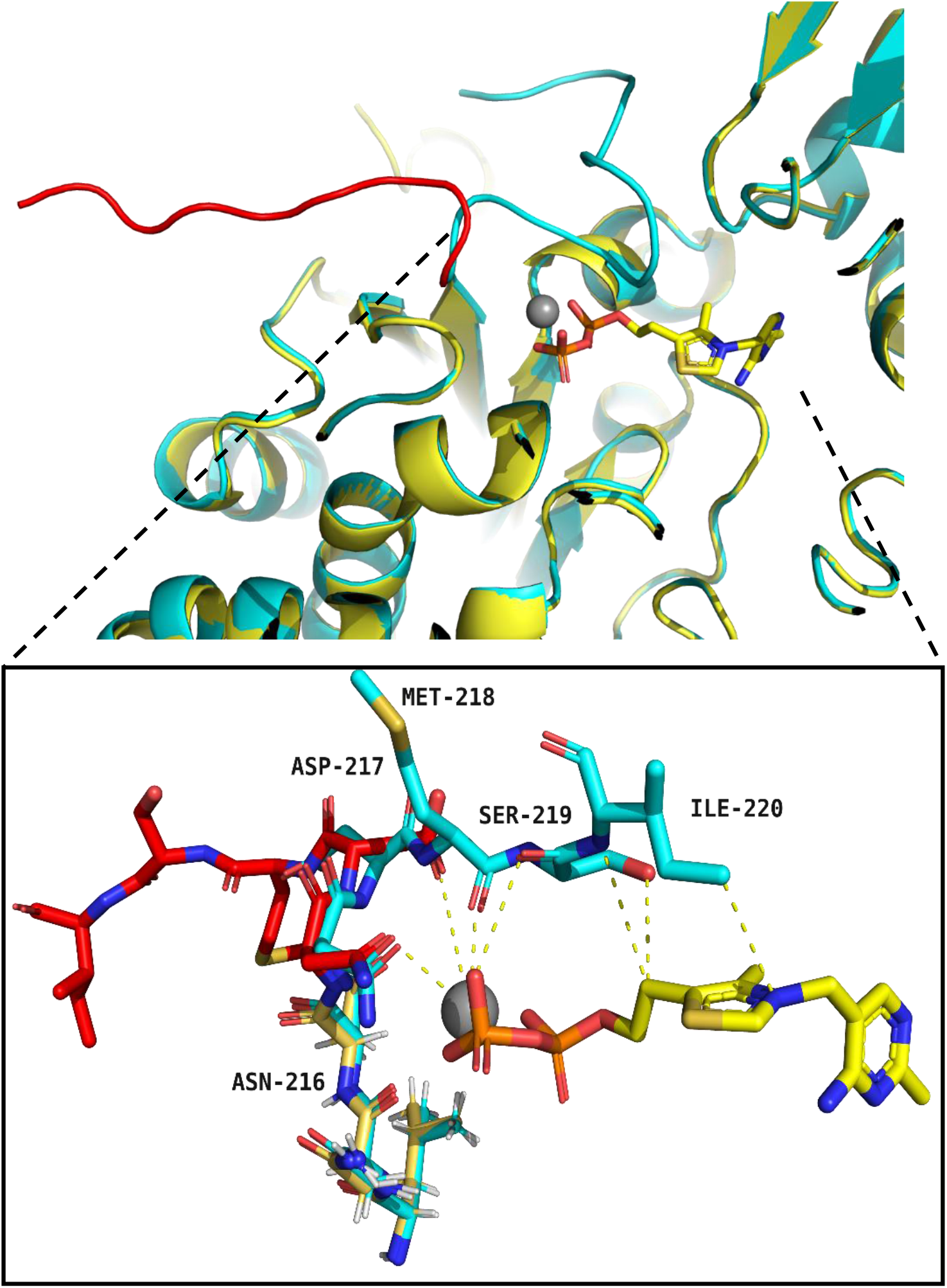
The structure of paDXPS co-crystallized with ThDP in cyan (PDB code: **8A5K**) is superimposed with the structure of apo paDXPS in yellow (PDB code: **8A29**), showing the conformational change in a loop close to the active site, RMSD 0.301 A° (6282 to 6282 atoms). Amino acids from the loop involved in ThDP binding are labeled with 3-letter code. Amino acids from the apo loop showing the open conformation are highlighted in red. ThDP atoms are shown as sticks: (C, yellow; O, red; N, blue; P, orange), Magnesium cation is shown as gray sphere, residue atoms are shown as sticks (C, cyan; O, red; N, blue; P, orange; S, yellow). Interactions are shown as yellow, dashed lines (below 5Å).

The *K*m value of ThDP for paDXPS was determined to be 86.9 nM, while a value of 114 nM was observed for kpDXPS. No kinetics values for ThDP were reported for other homologues, which makes our values difficult to interpret. However, kpDXPS is showing slightly lower affinity for ThDP in comparison to paDXPS, which may be due to the different shaping of the binding site. Next, the influence of cofactor binding on the shaping of the active site was evaluated by obtaining structural information on the apo version of paDXPS.

### Structural analysis of an apo form of paDXPS

Apo paDXPS crystals were generated by treating the protein with concentrated ammonium acetate solution to remove residual ThDP. Subsequently, the protein was resuspended in storage buffer and concentrated to 15 mg/mL before setting up hanging drops with the known crystal condition. As before, crystals were observed and grew in 3–5 days to 200 μm at 18°C. A complete dataset was collected at SLS beamline X06DA, the structure was solved using molecular replacement and refined to 2.2 Å (PDB ID: 8A29). There are slight differences seen in the unit cell parameters between the apo and the ThDP-bound structure, the apo unit cell is slightly bigger in size (a,116.44, b,137.63, c,232.08) than the ThDP co-crystal (a,115.927, b,133.524, c,231.461), which can be explained by the more open conformation exhibited by the apo enzyme. The superimposition of the apo structure with the ThDP-bound structure revealed an overall RMSD of 0.301Å (6282 to 6282 atoms). Upon comparing the structure of apo paDXPS with the cofactor-bound version, we could identify a region that is present in two different conformations. The loop extending from Asn216 to Glu247 that is directly adjacent to the ThDP binding site appears to move away (or at least heavily distorted), showing nearly 180° conformational change upon ThDP binding. In the cofactor-bound version, this exact site appears to be stabilized by the coordination of the Asn216 side chain and the Met218 backbone with the Mg^2+^ cation, which is, in turn, coordinated by the diphosphate. The complex structure also shows participation of the Ile220 side chain, showing the potential to form an arene-H interaction with the thiazole ring (Fig. 4). Intriguingly, the observation of the conformational change in the apo structure and the aforementioned loop is also observed when the protein was co-crystalized with a thiamine analogue inhibitor lacking the diphosphate moiety, which is discussed later (Fig. 6). The different shaping of the active site triggered by an absence of ThDP has important implications for inhibitor design, as a different three-dimensional fold needs to be targeted.

**Figure 5:**
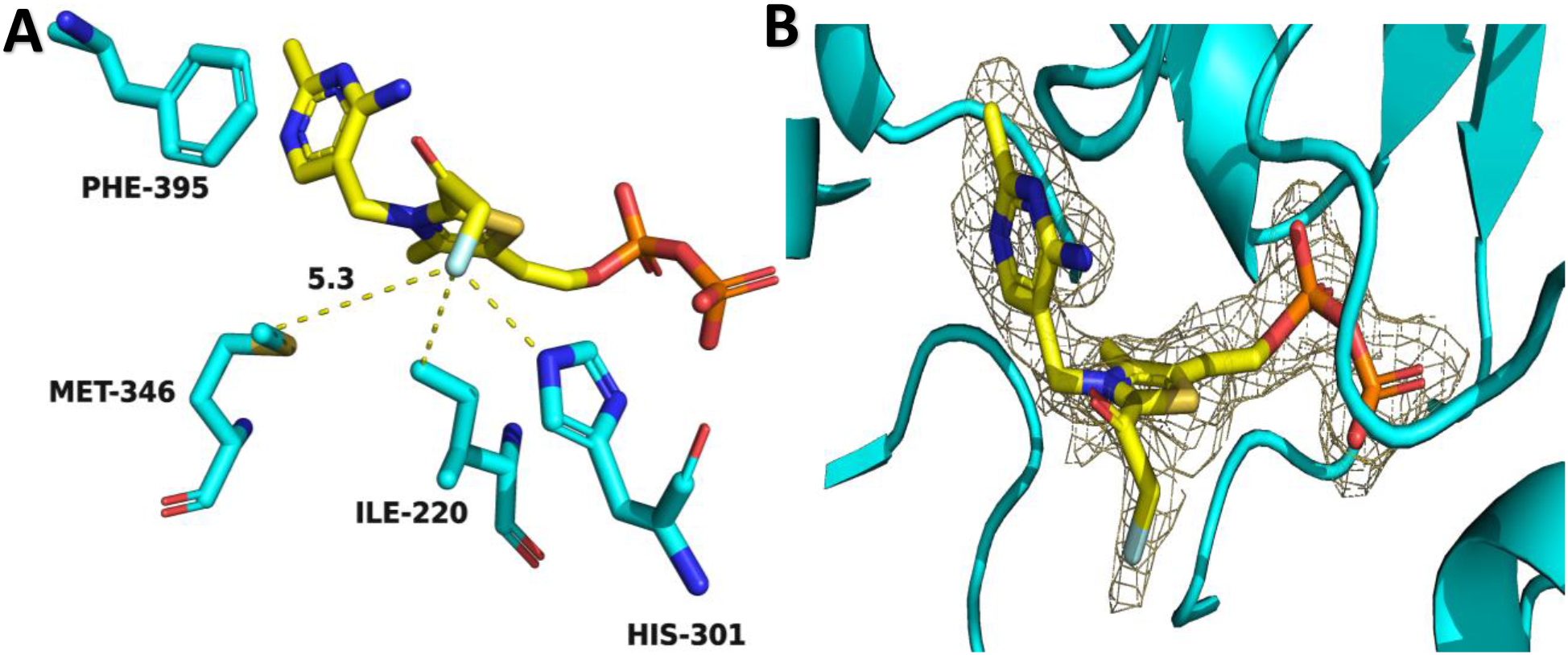
Binding of the 2-fluoroacetyl-ThDP with paDXPS (PDB ID: **8A45**). **A**: 2-fluoroacetyl-ThDP atoms are shown as sticks: (C, yellow; O, red; N, blue; P, orange; F, light green), residue atoms are shown as sticks (C, cyan; O, red; N, blue; P, orange; S, yellow). Interactions are shown as yellow dashed lines. **B:** Composite omit electron density maps of 2-fluoroacetyl-ThDP binding to paDXPS, the difference electron density map of the ligand (F_O_ - F_C_) was contoured at 3σ with phases calculated from a model that was refined in the absence of the ligand and is shown as a grey isomesh. Distances are given in Ångstrom.

**Figure 6:**
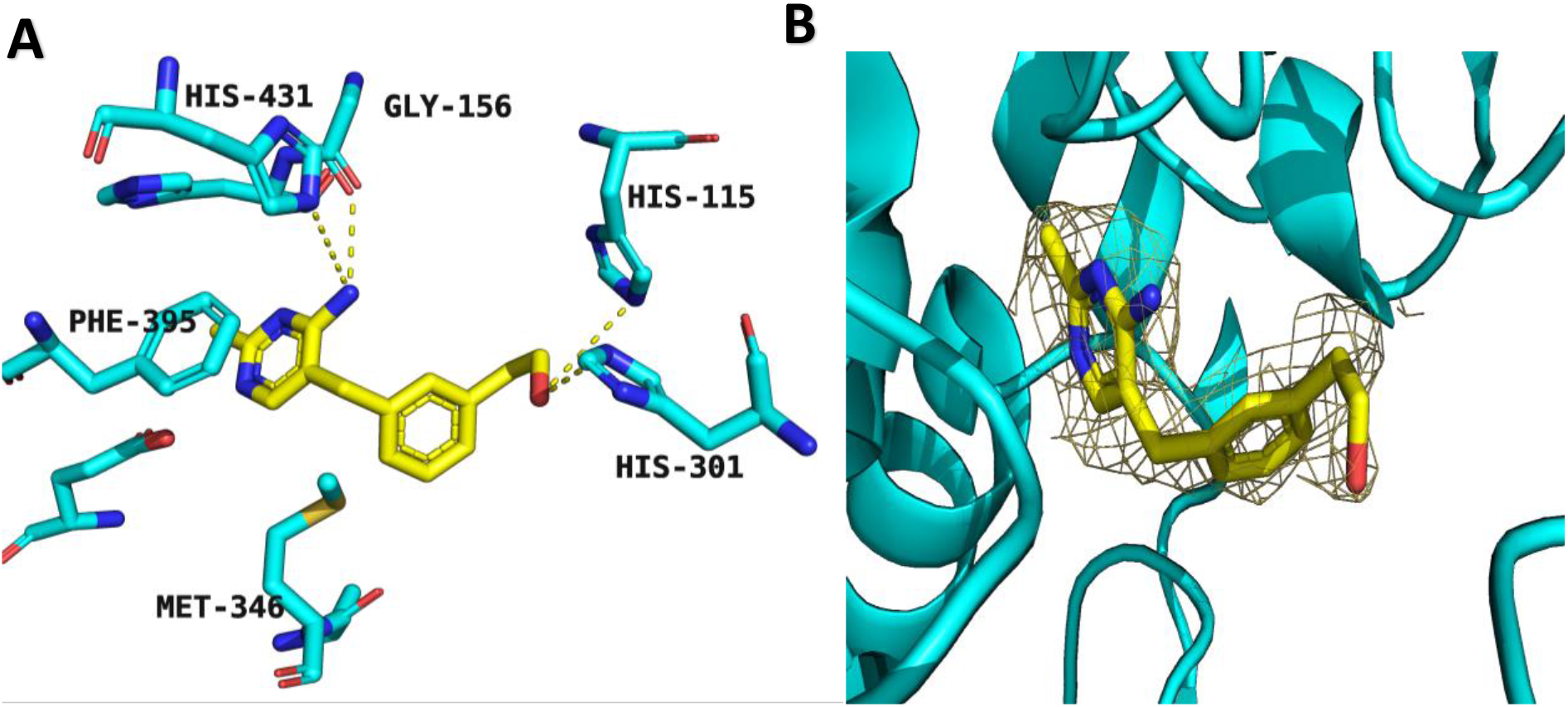
**A** Binding of thiamine analogue **1** with paDXPS (PDB ID: **8A4D**) (A). Interactions between thiamine analogue **1** and binding residues of *P. aeruginosa* DXPS. Thiamine analogue **1** atoms are shown as sticks: (C, yellow; O, red; N, blue), residue atoms are shown as sticks (C, cyan; O, red; N, blue). Interactions are shown as yellow, dashed lines (below 5Å). **B**: Composite omit electron density maps of inhibitor **(1)** to paDXPS, the difference electron density map of the ligand (F_O_ - F_C_) was contoured at 3σ with phases calculated from a model that was refined in the absence of the ligand and is shown as a grey isomesh.

### Co-crystallization of paDXPS with the substrate analogue fluoropyruvate

Fluoropyruvate is an analogue of the natural substrate pyruvate; it has been shown to deactivate the ThDP-dependent enzyme-pyruvate-dehydrogenase complex. The proposed mechanism of action has been reported to be the acetylation of a sulfhydryl group in the active site, which renders the enzyme inactive^7^. In addition, fluoropyruvate was also described as a first inhibitor of DXPS from *P. aeruginosa*^8^. In an *in vivo* situation, the ThDP-bound enzyme decarboxylates pyruvate upon interaction with ThDP to form the intermediate LThDP. This reaction promotes conformational changes, making the enzyme more receptive to DL-GAP ^11,19^.

In this study, we performed an inhibition assay and a mode-of-inhibition study on paDXPS using fluoropyruvate. In an inhibition assay, fluoropyruvate inhibits the enzyme paDXPS with a half-maximal inhibitory concentration (IC_50_) value of 80 μM. In the competitive assay, the IC_50_ value stayed in the same range (Table 2) despite the addition of up to 10-fold *K*_m_ of ThDP and pyruvate to the reaction mixture. In other words, paDXPS activity cannot be re-established by the cofactor and the natural substrate upon exposure to fluoropyruvate and the inhibition is non-competitive with both the cofactor and the substrate pyruvate^20^. An often-observed feature of non-competitive inhibitors is the binding to an allosteric site, which does not affect the catalytic pocket. Since fluoropyruvate is not expected to bind to an allosteric site and it has also been reported to inactivate the E1 subunit of the pyruvate dehydrogenase complex by covalently modifying a sulfhydryl group in the active site^7^, here we propose a similar reaction mechanism illustrating the interaction of fluoropyruvate with ThDP-bound paDXPS, depicted in scheme 1. To check whether the inhibition of the enzyme is the result of acetylation of a reactive residue in the active site after addition of fluroupyruvate, we performed native MS measurements to confirm acetylation of the enzyme by mass shift. We also conducted denatured MS analysis to detect covalent binding of one of reaction intermediates shown in scheme 1. In both cases, the MS results were not conclusive as we could not see distinct peaks neither corresponding to the apo protein nor the expected shift in mass, so we speculated that the protein is not tolerating the native MS conditions.

**Table 2:** MOI of fluoropyruvate: IC_50_ values calculated with varying concentration of ThDP and pyruvate (0.8–10-fold *K*_m_)

| [ThDP] (nM) | fluoropyruvate $IC_{50}$ ($\mu M$) | [Pyruvate] (mM) | fluoropyruvate $IC_{50}$ ($\mu M$) |
| --- | --- | --- | --- |
| 500 | $72.5 \pm 13$ | 1 | $83.7 \pm 8.7$ |
| 250 | $72.9 \pm 8$ | 0.5 | $81.2 \pm 27$ |
| 125 | $71.1 \pm 12$ | 0.25 | $79.2 \pm 26$ |
| 62.5 | $66.3 \pm 4$ | 0.125 | $51 \pm 1.6$ |
| 31.25 | $80.9 \pm 11$ | 0.0625 | $70 \pm 11$ |

Gathering structural information on a fluoropyruvate adduct of paDXPS would shed light on a covalently bound reaction product and deepen our knowledge about DXPS kinetics. For this, paDXPS (15 mg/mL) was incubated and crystallized with 0.5 mM fluoropyruvate in the presence of 0.5 mM ThDP and 5mM MgCl2. A complete dataset was collected at SLS beamline PXIII (X06DA), the structure was solved using molecular replacement and refined to 2.0 Å (PDB ID: 8A45).

We observed convincing electron density for the proposed reaction product (1), which will be referred to as (2-fluoro-acetyl ThDP) in the active site (Fig. 5). In the formerly mentioned report^21^, LThDP was shown to be stabilized by His434 and His51 in the reported drDXPS structure in complex with the enamine intermediate (corresponding to His431 and His84 in paDXPS). These histidine residues are also involved in stabilizing 2-fuoro-acetyl ThDP. Our structure also shows involvement of Ile220 (arene-H interaction) and His301 (salt bridge, 3.5 Å distance) in coordinating the thiazole and fluorine, respectively. This histidine residue (His301) is conservatively mutated to tyrosine in EcDXPS.

**Scheme 1:**
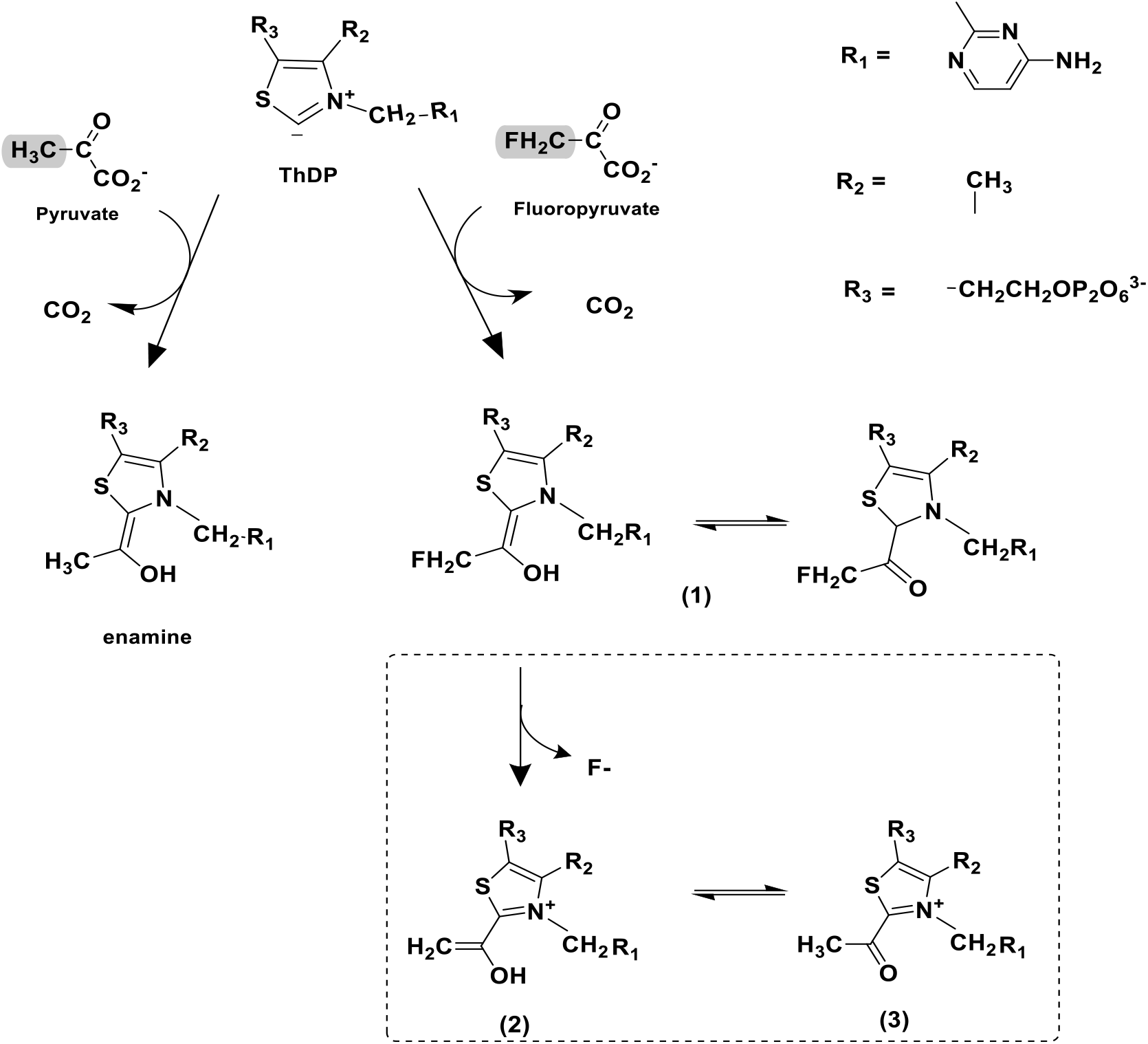
Suggested reaction mechanism of fluoropyruvate adduct (2-fluoro-acetyl ThDP) catalyzed by paDXPS in combination with ThDP in comparison to the reported reaction with the natural substrate pyruvate. The point of difference between fluoropyruvate and pyruvate is highlighted in gray.

### Co-crystallization of paDXPS with thiamine analogues

The thiamine analogue 3-deazathiamine diphosphate is the most potent reported inhibitor of several ThDP-dependent enzymes^22,23^ and several thiamine analogues were synthesized previously in our group to target DXPS as inhibitors^13,23^. In this study, we tested some of these compounds on DXPS from *P. aeruginosa* and *K. pneumoniae*. IC_50_ values are shown in Table 3, with the most potent compound **1** having an IC_50_ value of 73.7 ± 5.4 μM for kpDXPS. Subsequently, we set out to obtain an X-ray structure of either paDXPS or kpDXPS in complex with the thiamine analogues. We were unsuccessful to generate complex crystals for kpDXPS with any of the inhibitors. However, we were able to obtain a co-crystal structure of paDXPS with **1** by incubating paDXPS with 500 μM of **1** for 2 h on ice before setting up crystal plates (Fig. 6A). A dataset was collected at the SLS beamline X06SA, the structure was solved using molecular replacement and refined to 2.2 Å (PDB ID: 8A4D).

**Table 3:**
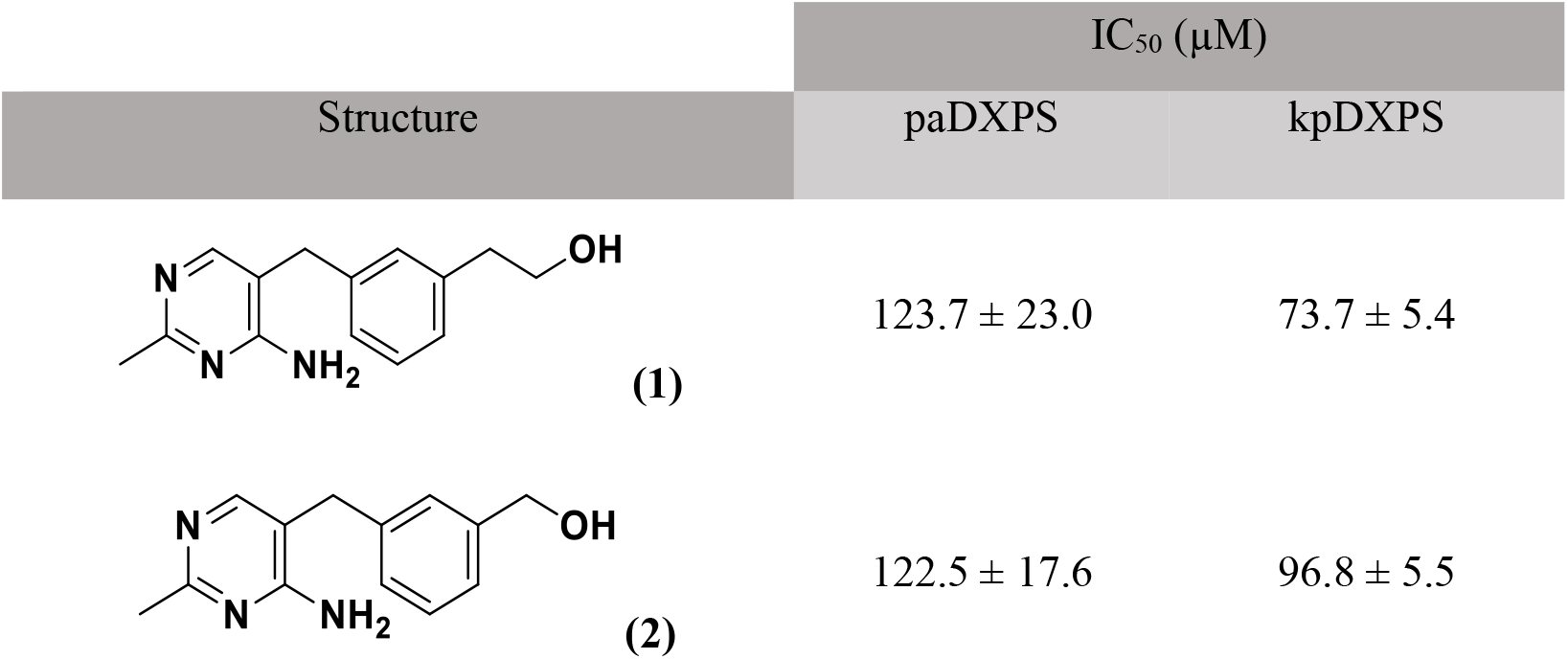
Biochemical evaluation of thiamine analogues as inhibitors of DXPS. The IC_50_ values on paDXPS and kpDXPS are shown.

| Structure | IC<sub>50</sub> (μM) |  |
| --- | --- | --- |
|  | paDXPS | kpDXPS |
| (1) | 123.7 ± 23.0 | 73.7 ± 5.4 |
| (2) | 122.5 ± 17.6 | 96.8 ± 5.5 |

As expected, binding of this analogue is similar to binding of the thiamine part (pyrimidine and thiazole) of ThDP with the pyrimidine ring stacking against Phe395. However, the phenyl ring of the inhibitor is showing a slight deviation from the plane of thiazole binding seen with ThDP and the hydroxyl group of the ligand is pointing towards the entry site of the substrates. Interestingly, binding of this inhibitor excludes parts of Domain I that are involved in coordination with the diphosphate in ThDP and this loop is showing the same conformation as seen in the apo enzyme (Figure 4), which adds to the notion that this loop is stabilized by the interaction of this specific part of ThDP.

### Structural analysis of apo and cofactor-bound kpDXPS

In addition to paDXPS, we also attempted to obtain structural information on another important homologue of pathogenic bacteria, namely DXPS from *Klebsiella pneumoniae*. Similar to paDXPS, we chose an optimized construct, in which we replaced the flexible loop (amino acids 197–240 of complete sequence) with six glycine residues, for crystallization because of an expected increase in protein stability. Crystallization trials for a cofactor-bound version of kpDXPS were successful using the sitting-drop vapour-diffusion method at a protein concentration of 13 mg/mL incubated together with 1 mM ThDP/MgCl_2_ and a reservoir solution consisting of 100 mM HEPES pH 7.0–7.5 and 18–28% PEG-3000. Plate-like crystals grew to 200 μm at 18 °C in about 20 days, belonged to space group P 2_1_ and diffracted to a maximum of 1.8 Å. A complete dataset was collected at SLS beamline X06SA located at the Swiss Light Source, and the structure was solved using molecular replacement with the published DXPS structure of *E. coli* (PDB ID: 2o1s).

There are two protein chains in the asymmetric unit, which form the expected biological dimer known from other DXPS enzymes. In contrast to paDXPS, this observation could not be backed up by native MS measurements due to solubility issues of kpDXPS. Instead, a stable assembly of the dimer could be confirmed by analyzing the structure with the PISA server^24^, revealing a buried interface area of 8820 Å^2^ and a free dissociation energy of Δ*G*_diss_ = 59.8 kcal/mol. Even though we determined a comparable affinity for ThDP, we can only observe convincing cofactor electron density in the B chain of kpDXPS (Fig. 7A). The three-dimensional shaping of the active site is similar to other homologues, with only one active site mutation (Ser52) towards the diphosphate-binding loop. Some crucial interactions for cofactor binding include ionic interactions of the pyrimidine with Gly121 and Glu336 as well as diphosphate coordination by Asn179 and Asn181 (Fig. 7B).

**Figure 7:**
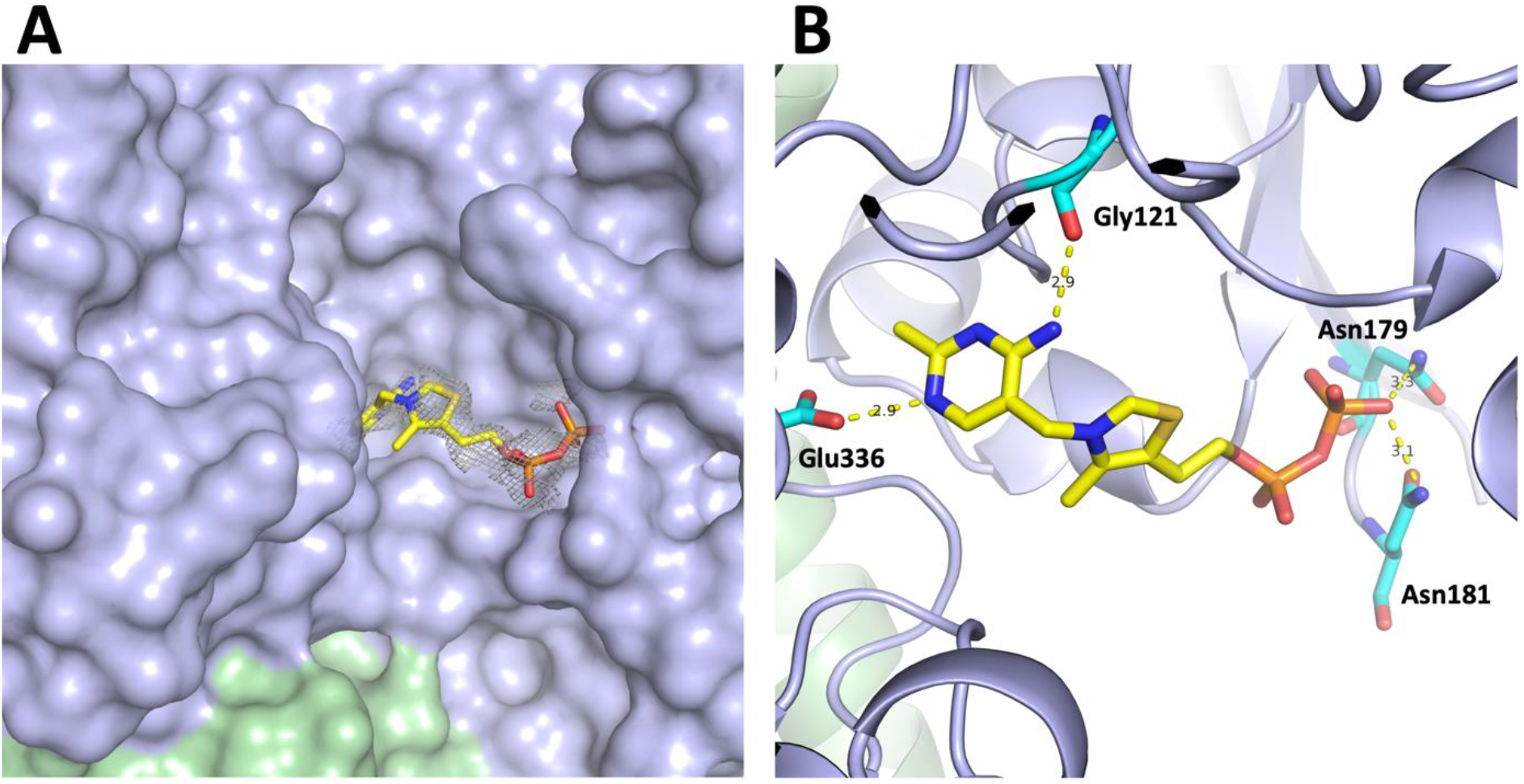
**A** Surface view of the kpDXPS (PDB ID: **8A9C**) active site. The two kpDXPS chains are depicted in light blue and palegreen, ThDP is shown as sticks with color-coded atoms (C: yellow, N: blue, O: red, S: ochre, P: orange). The difference electron density map of ThDP (F_O_ - F_C_) was contoured at 3σ with phases calculated from a model that was refined in the absence of ThDP and is shown as a grey isomesh. **B** Coordination of ThDP in kpDXPS. Interacting residues are shown as cyan sticks with color-coded atoms (C: cyan, N: blue, O: red). Distances are given in Ångstrom.

Intriguingly, we were unable to observe any crystals in our high-resolution crystal condition in the absence of ThDP addition, prompting us to search for a different condition. Crystals of apo kpDXPS were obtained at a considerably higher concentration than the cofactor-bound version at 47.5 mg/mL with a reservoir solution containing 200–400 mM MgCl_2_, 100 mM HEPES pH 7.5 and 14–18% PEG-8000 and grew within 5 days at 18 °C up to 300 μm. They belong to space group C 2221, diffracted to about 2.1 Å and a complete dataset was collected at ESRF beamline ID23-1. Subsequently, the structure was solved using molecular replacement with a refined ThDP-bound kpDXPS model.

As for the cofactor-bound structure, we found two protein chains in the asymmetric unit, which form the expected biological dimer. Both structures superimpose with an RMSD of 0.336 Å over 5514 atoms, which already suggests considerable movement between the two structures. Indeed, we observe significant loop rearrangement in the apo structure around the active site, as three loops are extending more towards a now empty pocket with mean distances around 4Å (Fig. 8A). This is an especially critical observation for drug discovery, as a different three-dimensional shape of the active site could enable an exploration of different chemical space in search for DXPS inhibitors. In addition, a loop belonging to domain I of DXPS (amino acids 91–124) is completely disordered in the apo structure of kpDXPS, leading to a more open conformation towards the other site of the substrate entrance channel (pale green region in Fig. 8AB). This observation is also backed up by the surface representation (Fig.8B), which nicely depicts a narrower cofactor pocket and a tunnel opening up on the other side of ThDP due to the aforementioned disordered loop.

**Figure 8:**
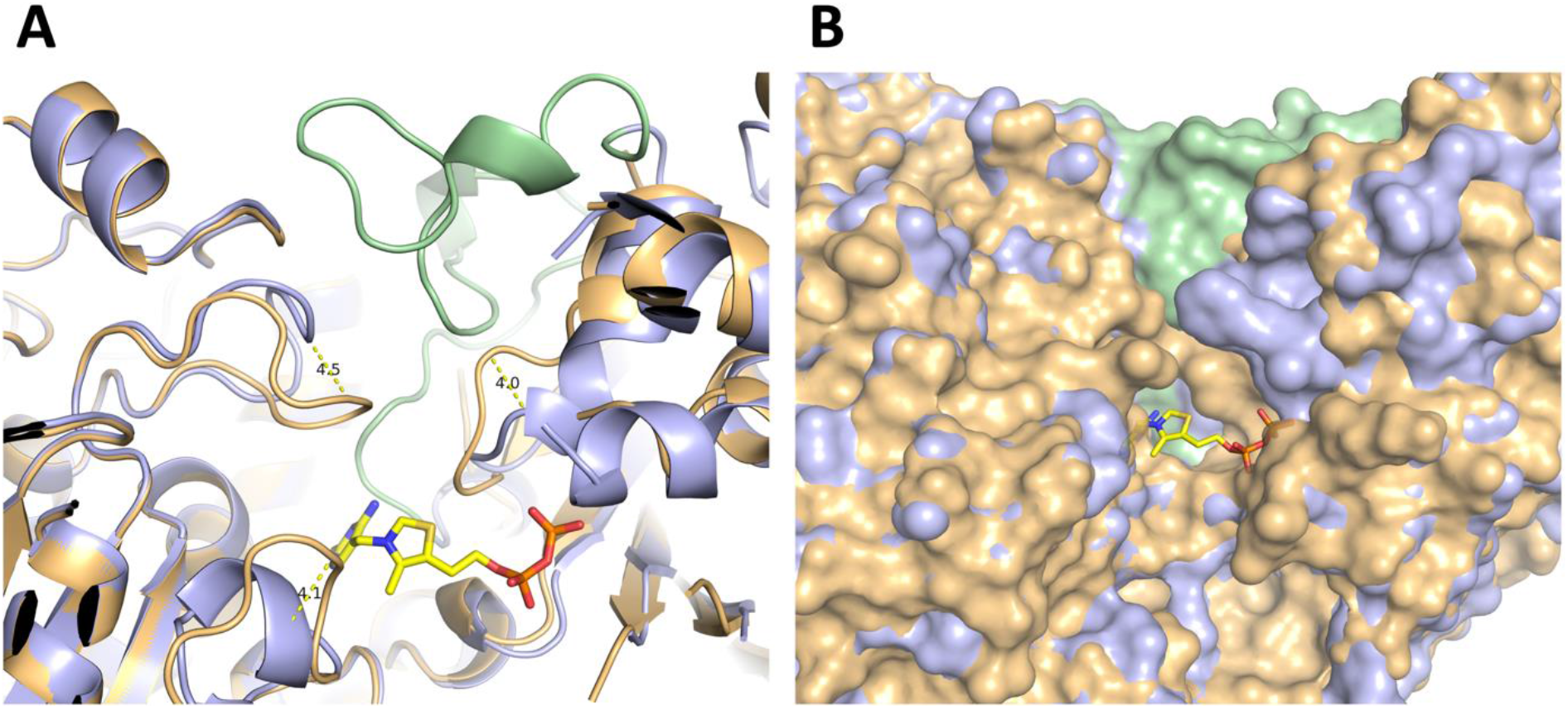
**A** Loop rearrangement close to the active site in apo kpDXPS (PDB ID: **8A8Y**). The two versions of kpDXPS are visualized as a cartoon representation in light blue (ThDP-bound) and light orange (apo). An N-terminal disordered loop in apo kpDXPS (amino acids 91–124) is shown in pale green. ThDP is shown as sticks with color-coded atoms (C: yellow, N: blue, O: red, S: ochre, P: orange). Distances between the loops are given in Ångstrom. **B** Comparison of the surface representation of the kpDXPS active site, with the same color-coding as in A. Loop movement leads to a much narrower pocket in apo kpDXPS, however the missing N-terminal loop opens up a tunnel on the other side of the cofactor-binding site.

### Structural comparison of paDXPS and kpDXPS

KpDXPS and paDXPS share a sequence identity of 62.5% and similarity of 84.3% (Blosum score ≥ 0). Predictably, the structural arrangement is also highly similar, the two structures superimpose with an RMSD of only 0.663 Å (4797 to 4797 atoms) when comparing both ThDP-bound crystal structures (Fig. 9). kpDXPS is also showing the shorter B-sheet in domain I (residue 215–217) seen in paDXPS, however, the loops in the kpDXPS structure arrange slightly different. Moreover, kpDXPS is missing residues (246–248) due to lack of density, which are part of the loop connecting domains I and II and are also part of the so-called spoon-fork motif. The active site of kpDXPS shows an almost identical binding mode of ThDP and similar residues are involved in binding, except for Ser52, which is mutated to alanine in the paDXPS structure. This residue might be an important distinction for drug design, as the hydroxyl group of Ser52 adds a targetable nucleophilic moiety to the active site. Furthermore, the active site of kpDXPS shows exclusion of a histidine residue from the active site when compared to the corresponding histidine (His115) in paDXPS. His80 in kpDXPS moves out of the active site by a deviation of 5.5Å, showing a contact distance of 8.7Å versus 2.1Å for HIS115 in paDXPS. It is unclear to us whether removal of this positively charged residue from the binding site is the reason why kpDXPS has less affinity to ThDP and therefore shows increased inhibition by ThDP analogues in general (table 2). Unfortunately, there is not enough density for the diphosphate-binding loop in the kpDXPS structure to compare with the one from paDXPS. Intriguingly, loops near the active site in kpDXPS seem to undergo a different conformational change, which results in the aforementioned differences in three-dimensional shape of the pocket (Fig. 8).

**Figure 9:**
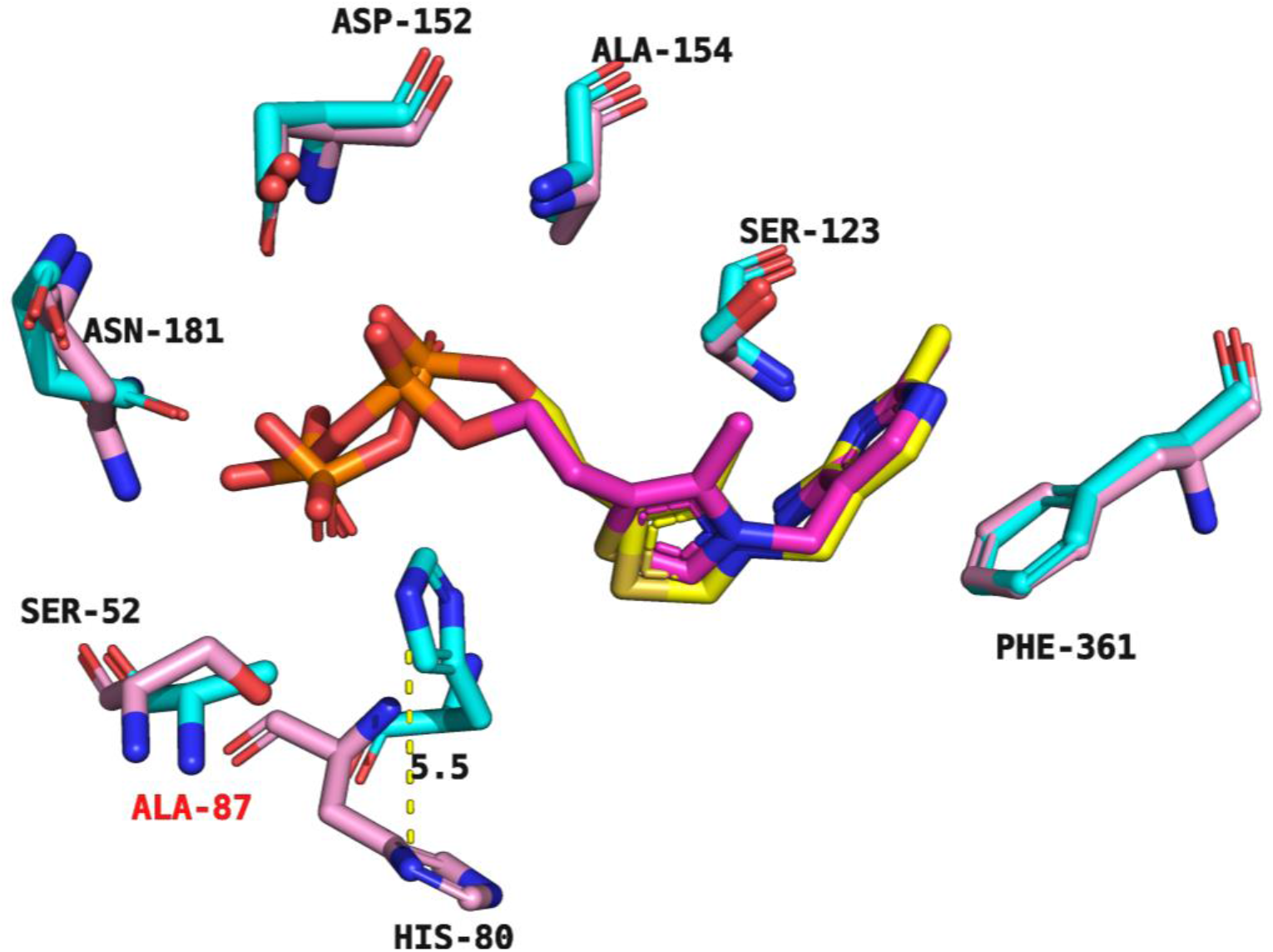
Comparison of ThDP-bound kpDXPS and paDXPS. Residues shaping the active site and involved in binding to ThDP are highlighting differences in the active site. PaDXPS is shown in cyan, overlaid with kpDXPS in pink, a mutated residue in paDXPS is labeled in red (Ala to Ser52). ThDP atoms from paDXPS are shown as sticks: (C, yellow; O, red; N, blue; P, orange), ThDP atoms from kpDXPS are shown as sticks: (C, magenta; O, red; N, blue; P, orange). Residue atoms are shown as sticks (C, cyan; O, red; N, blue; P, orange; S, yellow). Distance change between His80 in kpDXPS and the aligned histidine in paDXPS is shown as yellow, dashed lines.

## Conclusions

The structures of *P. aeruginosa* and *K. pneumoniae* DXPS alone and in complex with the cofactor ThDP broaden the knowledge about this important class of enzymes and provide valuable structural information for the development of new drugs targeting ESKAPE pathogens that may eventually be used to treat multidrug-resistant infections.

From a drug-design perspective, these results open the door to different modes of DXPS inhibition. Our structures reveal the important role of the diphosphate moiety of ThDP in stabilizing part of the active site. Targeting specific residues in this loop may prevent the enzyme from binding to ThDP by not providing enough space for ThDP to bind, especially with the rearrangement of the loops in kpDXPS. In paDXPS, one strategy could be to prevent the diphosphate binding loop from closing, thereby preventing the enzyme from entering its active conformation. This could perhaps be achieved by using the pyrimidyl ring present in ThDP as a warhead and optimizing the rest of the compound to bind to the loop in its “open” conformation. Additionally, the binding mode of thiamine analog **1** indicates that it is also possible to explore compounds that target the substrate binding channel in addition to the ThDP binding pocket. Following this approach, a three-headed inhibitor could perhaps be developed, which would likely show increased selectivity for DXPS over other ThDP-dependent enzymes due to the exploitation of DXPS’s unique binding-site geometry. Furthermore, we demonstrated that a non-competitive inhibition could be achieved by blocking further interactions of ThDP-bound enzyme with molecules that can form direct intermediates with ThDP, therefore trapping the enzyme in this rate-determining kinetic step and preventing subsequent interactions with the natural substrates. These additional DXPS X-ray crystal structures from important ESKAPE pathogens lay the foundation for future structure-based drug design of potent and selective small-molecule inhibitors to treat infections worldwide.

## Materials and methods

### Protein Expression and Purification

#### paDXPS

The construct genes were synthesized commercially, cloned into pET28a and transformed into BL21(DE3) for overexpression. The transformed cells were grown in a selective LB medium supplemented with Kanamycin (50 mg/mL) up to an OD600 of 0.6, then induced with 1 mM isopropyl-L-D-thiogalactoside (IPTG) and were subsequently grown at 18 °C for 16 h. The harvested cells were then disrupted in a microfluidizer after resuspension in washing buffer consisting of 50 mM HEPES, 100 mM NaCl, 20 mM imidazole and 2 mM β-mercaptoethanol (100 mL buffer per 25 g of wet cell pellet). After centrifugation, the supernatant was loaded onto a 5 mL HisTrap HP column equilibrated in washing buffer. After an extensive washing step with 30 column volumes of washing buffer, the bound fraction was then eluted with 50 mM HEPES, 100 mM NaCl, 300 mM Imidazole and 2 mM β-mercaptoethanol. His-tag purification was followed by size-exclusion chromatography in 20 mM HEPES, 200 mM NaCl and 2 mM β-mercaptoethanol. Sodium dodecyl sulfate polyacrylamide gel electrophoresis (SDS-PAGE) and protein mass spectrometry was used to examine the purity of protein purifications.

#### kpDXPS

The construct gene was synthesized commercially, cloned into pET28b+ and transformed into Lemo21(DE3) for overexpression. The transformed cells were grown in a selective LB medium at 37 °C supplemented with kanamycin (50 mg/mL) and chloramphenicol (34 mg/mL) up to an OD_600_ of 0.9, then induced with 0.1 mM isopropyl-L-D-thiogalactoside (IPTG) and were subsequently grown at 18 °C for 16 h. The harvested cells were then disrupted in a microfluidizer after resuspension in washing buffer consisting of 20 mM Tris pH 8.0, 200 mM NaCl, 20 mM Imidazole pH 8.0 and 5% glycerol (100 mL buffer per 25 g of wet cell pellet). After centrifugation, the supernatant was loaded onto a 5 mL HisTrap HP column equilibrated in washing buffer. After an extensive washing step with 30 column volumes of washing buffer, the bound fraction was then eluted with an elution buffer consisting of 20 mM Tris pH 8.0, 200 mM NaCl, 250 mM imidazole pH 8.0 and 5% glycerol. Eluted protein was passed over a Hiprep 26/10 desalting column equilibrated in washing buffer to remove excess imidazole, after which TEV digestion was carried out by adding TEV protease in a 1:10 protein: TEV ratio and incubating the solution at 4°C for 16 h. The N-terminal 6x His-Tag was removed by passing the protein over a 5 mL HisTrap HP column and collection of the flow-through. Purification was finalized by size-exclusion chromatography in 20 mM Tris, 200 mM NaCl, final pH 8.0 for apo kpDXPS and 10 mM HEPES, 200 mM NaCl, 5% glycerol, final pH 7.4 for ThDP-bound kpDXPS. Sodium dodecyl sulfate polyacrylamide gel electrophoresis (SDS-PAGE) and protein mass spectrometry were used to examine the purity of protein purifications.

### Crystallization

#### paDXPS

Crystallization trials were carried out with a protein solution containing 15 mg/mL paDXPS in storage buffer using commercially available screens from NeXtalbiotech. Complex structures were obtained after incubating paDXPS under the specific conditions mentioned in the respective sections. In every case, crystals were grown in hanging drops with a reservoir solution consisting of 100 mM HEPES, 12% PEG-8000 and 200 mM calcium acetate. Plates were observed every three days and optimization of the identified conditions was carried out. 32% Glycerol was added as a cryo-protectant before the crystals were mounted in a cryo-loop and stored in liquid nitrogen until data collection.

#### kpDXPS

Crystallization trials were carried out with a protein solution containing 12.5 mg/mL kpDXPS (complex) and 47.5 mg/mL kpDXPS (apo) in the respective storage buffer using commercially available screens from NeXtalbiotech. Complex structures of kpDXPS were obtained by addition of 1 mM ThDP/MgCl2 and incubating the solution for 4 h on ice before setting up the crystallization plates. For the protein complex, crystals were grown in sitting-drop optimization plates at 18 °C with a reservoir solution consisting of 100 mM HEPES pH 7.0–7.5 and 18–28% PEG-3000. For the apo version, crystals were grown in hanging-drop optimization plates at 18 °C with a reservoir solution consisting of 200–400 mM MgCl2, 100 mM HEPES pH 7.5 and 14-18% PEG-8000. In both cases, 15% 2*R*,3*R*-butanediol was added as a cryoprotectant before the crystals were mounted in a cryoloop and stored in liquid nitrogen until data collection.

### Data collection and refinement

X-ray diffraction data for paDXPS structures were collected at the Swiss Light Source located at the Paul Scherrer Institute in Switzerland. Specific information about each beamline can be found in the specific sections, all data collection and refinement statistics can be found in Table S2. Data reduction and scaling were done using AIMLESS^25^ in CCP4i ^26^. Structures were solved using molecular replacement using the program Phaser.MR ^27^ in Phenix software ^28^. Repetitive cycles of refinement were done in COOT^29^ and Phenix.refine^30^ to obtain final structures (table S1 and 2). Structures were validated using the Molprobity server ^31^ and all figures were rendered using PyMoL^32^.

### Kinetics and inhibition assays

To measure DXPS activity and kinetic parameters, a coupled spectrophotometric enzyme assay was adapted from the assay protocol as described by Altincicek *et al*.^8^. For inhibition studies, a reaction mixture A (100mM HEPES pH 7.5, 100mM NaCl, 400nM ThDP, 1mM MgCl2, 0.5μM NAPDH, 1μM IspC and 0.2μM paDXPS) was pre-incubated with different concentrations of inhibitor for 5 min at 37 °C in 10% DMSO, the reaction was then started by adding mixture B (100mM HEPES pH 7.5, 100mM NaCl, 2mM DL-glyceraldehyde-3-phosphate and 2mM pyruvate).

## Acknowledgements

The authors would like to thank Nicolas Frank for native MS and intact mass MS measurements. We acknowledge the Paul Scherrer Institute, Villigen, Switzerland for provision of synchrotron radiation beamtime at beamline X06DA-PXIII and X06SA-PXI of the SLS and the European Synchrotron Radiation Facility (ESRF), Grenoble, France for provision of beamline ID23-1.

## Author contributions

A.K.H.H., R.H. and S.A. conceived the study. R.H., S.A. wrote the main manuscript text and prepared all figures. A.L. and L.M. synthesized the compounds for bioassays. All authors proofread the manuscript. A.K.H.H supervised the project.

