## Supplemental for "Structural analysis of 1-deoxy-D-xylulose 5-phosphate synthase from *Pseudomonas aeruginosa* and *Klebsiella pneumoniae* reveals conformational changes upon cofactor binding"

**2. Department of Pharmacy, Saarland University, 66123 Saarbrucken.**

**3- Stratingh Institute for Chemistry, University of Groningen, Nijenborgh 7, NL-9747 AG Groningen, the Netherlands.**

**Table S1**: Primary amino acid sequences of proteins used in this study.

| **Construct** | **Primary amino acid sequence** |
|---|---|
| Native paDXS | MGSSHHHHHHSSGLVPRGSMENLYFQSHMPKTLHEIPRERPATPLLDRASSPAELRRLGEADLETLADELRQYLL YTVGQTGGHFGAGLGVVELTIALHYVFDTPDDRLVWDVGHQAYPHKILTERRELMGTLRQKNGLAAFPRRAESE YDTFGVGHSSTSISAALGMAIAARLQGKERKSVAVIGDGALTAGMAFEALNHASEVDADMLVILNDNDMSISHN VGGLSNYLAKILSSRTYSSMREGSKKVLSRLPGAWEIARRTEEYAKGMLVPGTLFEELGWNYIGPIDGHDLPTLVA TLRNMRDMKGPQFLHVVTKKGKGFAPAELDPIGYHAITKLEAPGSAPKKTGGPKYSSVFGQWLCDMAAQDARL LGITPAMKEGSDLVAFSERYPERYFDVAIAEQHAVTLAAGMACEGMKPVVAIYSTFLQRAYDQLIHDVAVQHLD VLFAIDRAGLVGEDGPTHAGSFDISYLRCIPGMLVMTPSDEDELRKLLTTGYLFDGPAAVRYPRGSGPNHPIDPDL QPVEIGKGVVRRRGGRVALLVFGVQLAEAMKVAESLDATVVDMRFVKPLDEALVRELAGSHELLVTIEENAVMG GAGSAVGEFLASEGLEVPLLQLGLPDYYVEHAKPSEMLAECGLDAAGIEKAVRQRLDRQ |
| Mutated paDXS | MGSSHHHHHHSSGLVPRGSMENLYFQSHMPKTLHEIPRERPATPLLDRASSPAELRRLGEADLETLADELRQYLL YTVGQTGGHFGAGLGVVELTIALHYVFDTPDDRLVWDVGHQAYPHKILTERRELMGTLRQKNGLAAFPRRAESE YDTFGVGHSSTSISAALGMAIAARLQGKERKSVAVIGDGALTAGMAFEALNHASEVDADMLVILNDNDMSISHN VGGLSNYLAKIGGGGGGPGTLFEELGWNYIGPIDGHDLPTLVATLRNMRDMKGPQFLHVVTKKGKGFAPAELD PIGYHAITKLEAPGSAPKKTGGPKYSSVFGQWLCDMAAQDARLLGITPAMKEGSDLVAFSERYPERYFDVAIAEQ HAVTLAAGMACEGMKPVVAIYSTFLQRAYDQLIHDVAVQHLDVLFAIDRAGLVGEDGPTHAGSFDISYLRCIPG MLVMTPSDEDELRKLLTTGYLFDGPAAVRYPRGSGPNHPIDPDLQPVEIGKGVVRRRGGRVALLVFGVQLAEAM KVAESLDATVVDMRFVKPLDEALVRELAGSHELLVTIEENAVMGGAGSAVGEFLASEGLEVPLLQLGLPDYYVEH AKPSEMLAECGLDAAGIEKAVRQRLDRQ |
| Mutated kpDXS | MSFDIAKYPTLALVDSTQELRLLPKESLPKLCDELRRYLLDSVSRSSGHFASGLGTVELTVALHYVYNTPFDRLIWDV GHQAYPHKILTGRRDKIGTIRQKGGLHPFPWRGESEYDVLSVGHSSTSISAGIGVAIAAAKEDKQRRAVCVIGDGA ITAGMAFEAMNHAGDIKPDLLVVLNDNEMSISENVGALNNHLAGGGGGGPGTLFEELGFNYIGPVDGHDVLG LVSTLKNMRDLKGPQFLHIMTKKGRGYEPAEKDPITFHAVPKFDHTSGVLPKSSGGLPSYSKIFGDWLCETAAKD NKLMAITPAMREGSGMVEFSKKFPDRYFDVAIAEQHAVTFAAGLAIGDYKPVVAIYSTFLQRAYDQVIHDVAIQK LPVLFAIDRAGIVGADGQTHQGAFDLSFLRCIPDMVVMTPSDENECRQMLYTGYHYSDGPCAVRYPRGSGTGA TLEPLASLPIGKGVVKRQGEKIAILNFGTLLPEAAAVADKLNATLVDMRFVKPLDTALILQLAGEHDALVTLEENAI MGGAGSGVNEVLMAHRRAVPVLNIGLPDYFIPQGTQEEIRADLGLDAAGIEAKIRDWLA |

**Table S2**: Data collection and refinement statistics of all X-ray structures shown in the manuscript. Statistics for the highest-resolution shell are shown in parentheses.

| | Apo paDXS | ThDP-bound paDXS | paDXS – 2-fluoroacetyl-ThDP | paDXS – Thaimine analog | ThDP-bound kpDXS | Apo kpDXS |
|---|---|---|---|---|---|---|
| PDB code | 8A29 | 8A5K | 8A45 | 8A4D | 8A9C | 8A8Y |
| Resolution range | 48.59–2.2 | 46.29–2.37 | 48.59–2.0 | 48.54–2.2 | 43.37 – 1.8 | 91.87 – 2.1 |
| Space group | P21 21 21 | P21 21 21 | P21 21 21 | P21 21 21 | P21 | C2 2 21 |
| Unit cell<br>a b c [Å]<br>α β γ [°] | 116.44 137.63 232.08<br>90.00 90.00 90.00 | 115.927 133.524 231.461<br>90 90 90 | 117.12 138.01 231.63<br>90.00 90.00 90.00 | 116.618 137.618 231.681<br>90.00 90.00 90.00 | 89.67 72.56 91.44<br>90.00 108.46 90.00 | 119.16 144.25 142.91<br>90.0 90.0 90.0 |
| Total reflections | 376678 (37313) | 901451 (46057) | 374926 (36518) | 371083 (36366) | 557529 (75397) | 323919 (45609) |
| Unique reflections | 188736 (18663) | 139116 (6871) | 187627 (18663) | 188257 (18546) | 100675 (14606) | 71565 (10344) |
| Multiplicity | 7.1 (7.3) | 6.5 (6.7) | 12.2 (12.3) | 4.5 (4.5) | 5.5 (5.2) | 4.5 (4.4) |
| Wavelength | 0.97 | 0.97 | 0.97 | 0.97 | 0.97 | 0.97 |
| Completeness (%) | 99.9 (100) | 99.8 | 99.62 (99.92) | 99.63 | 97.9 (97.6) | 99.6 (99.6) |
| Mean I/ sigma (I) | 11.09 (2.65) | 9.33 (2.08) | 18.01 (6.11) | 6.86 (1.65) | 13.4 (2.5) | 9.9 (2.2) |
| Wilson B-factor | 19.66 | 42.12 | 19.67 | 25.24 | 25.41 | 35.38 |
| R-merge | 0.140 (0.953) | 0.06354 (0.5769) | 0.150 (1.187) | 0.1092 (0.5661) | 0.066 (0.561) | 0.089 (0.720) |
| R-work | 0.1896 | 0.2650 | 0.1948 | 0.1996 | 0.1593 | 0.1749 |
| R-free | 0.2254 | 0.2950 | 0.2261 | 0.2323 | 0.1902 | 0.1987 |
| Number of non-hydrogen atoms | 27762 | 25544 | 28232 | 28005 | 8837 | 7781 |
| Macromolecules | 25575 | 25177 | 25744 | 25672 | 7984 | 7384 |
| Ligands | 70 | 172 | 296 | 213 | 26 | |
| Solvent | 2157 | 259 | 2300 | 2222 | 827 | 397 |
| Protein residues | 3357 | 3309 | 3384 | 3366 | 1043 | 970 |
| RMS (bonds) | 0.003 | 0.019 | 0.005 | 0.002 | 0.006 | 0.003 |
| RMS (angles) | 0.63 | 2.23 | 0.70 | 0.55 | 0.79 | 0.56 |
| Average B-factor | 25.78 | 57.10 | 29.70 | 31.57 | 32.6 | 48.29 |
| Macromolecules | 26.85 | 58.45 | 29.30 | 25672 | 31.84 | 48.42 |

*Statistics for the highest-resolution shell are shown in parentheses.

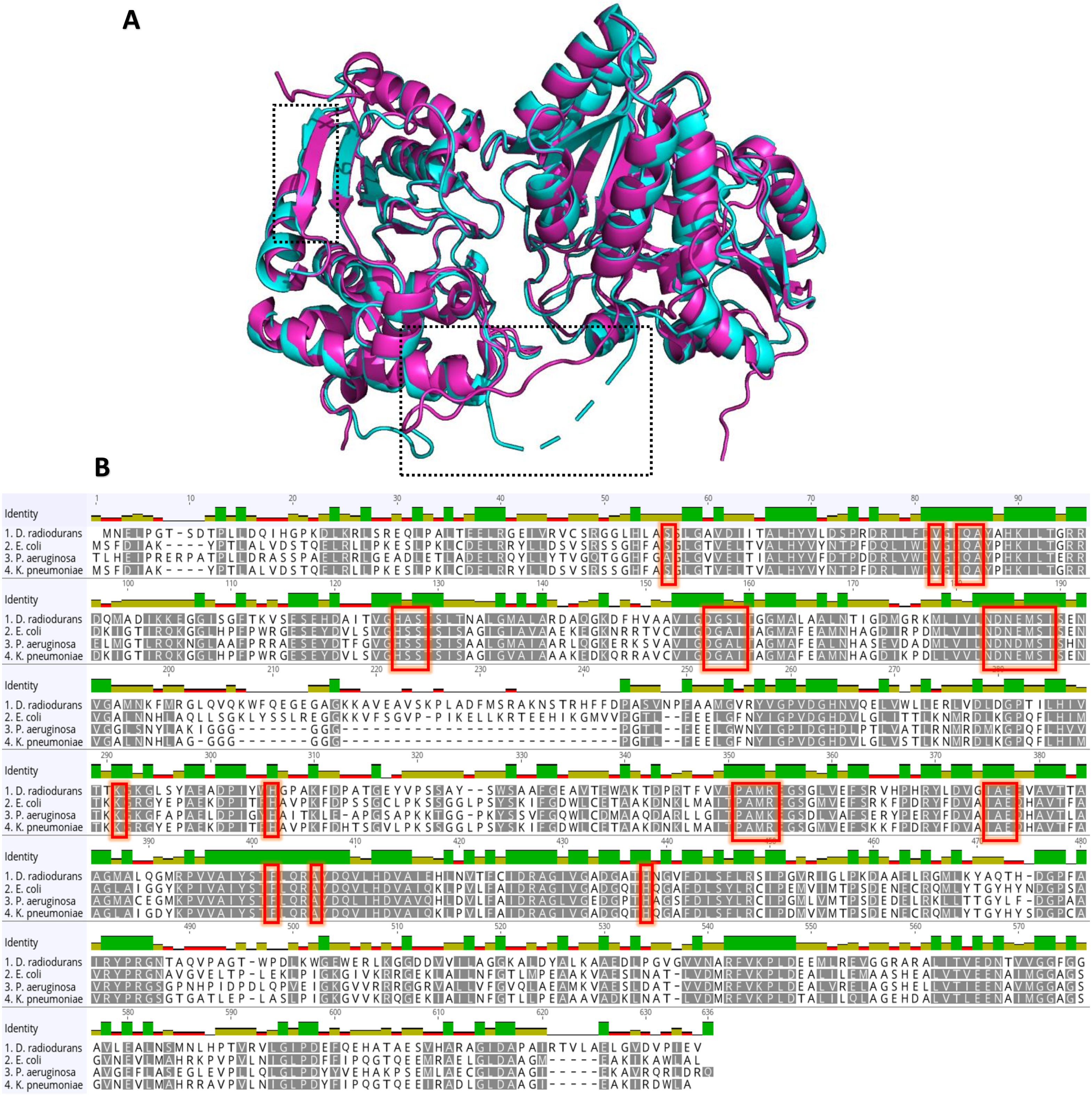


**Figure S1**: Comparison of paDXS and orthologues. **A:** Superimposition of paDXS (cyan) with drDXS 2O1X (light magenta), RMSD 0.778 Å, locations of difference are marked with dashed boxes. paDXS with e.coli DXS (2O1S ) (not shown) RMSD 0.913 A°. **B**: Sequence alignment of DXS from *p. aeruginosa*, *K. pneumonia*, *d. radiodurans* and *E.coli*. The identity, shown as bar graph above the sequences, was calculated using the four aligned sequences. Amino acids in the active site are indicated by red boxes.

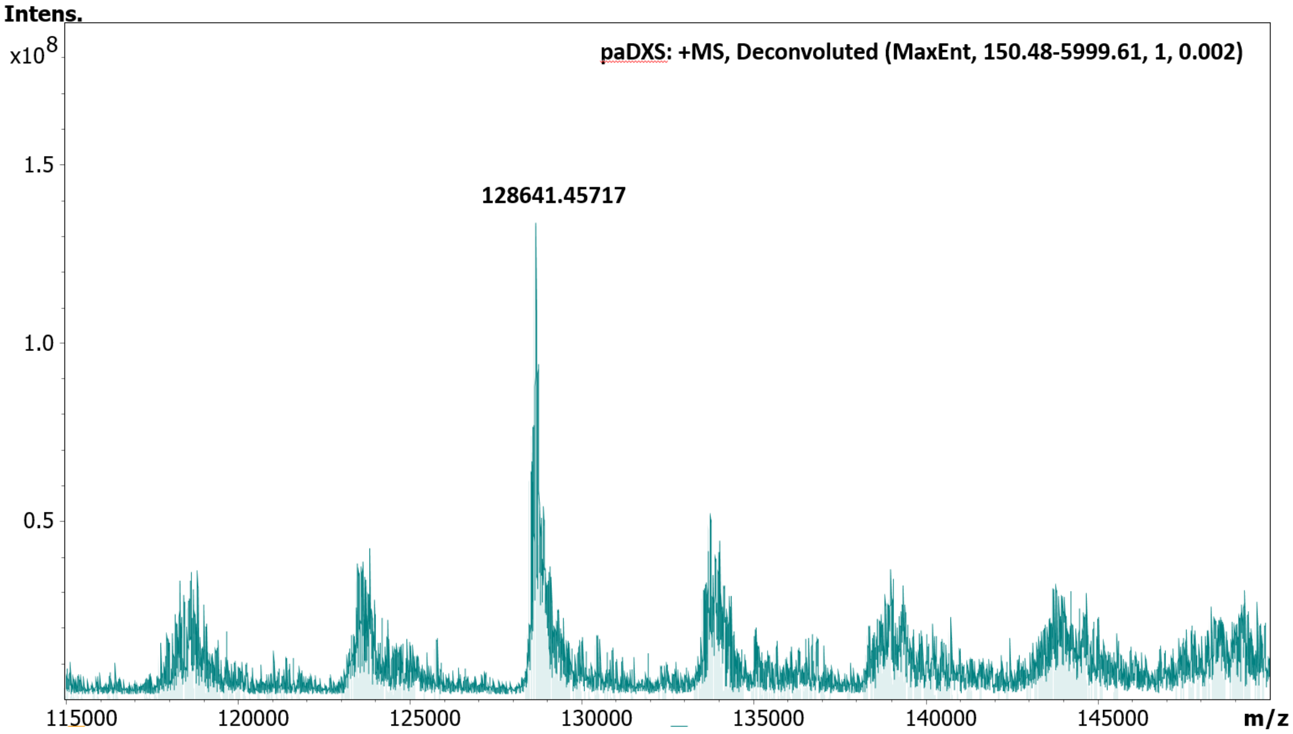


**Figure S2**: Native Ms analysis of paDXS. A peak at 128641 confirms the presence of paDXP as a dimer in solution.

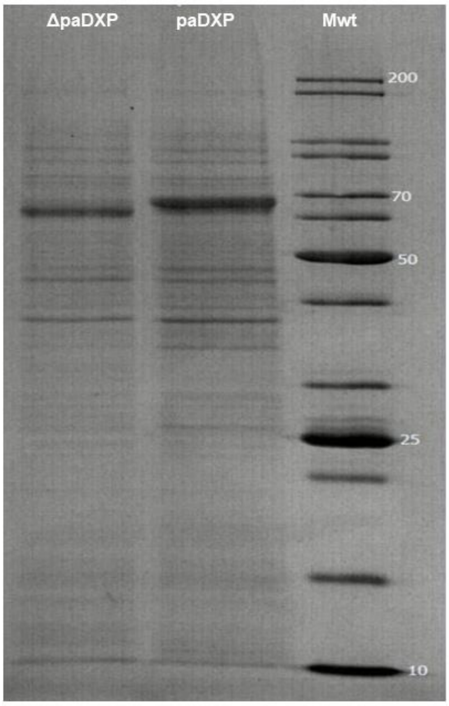


**Figure S3**: SDS-PAGE gel analysis of purified native and mutated paDXPS.

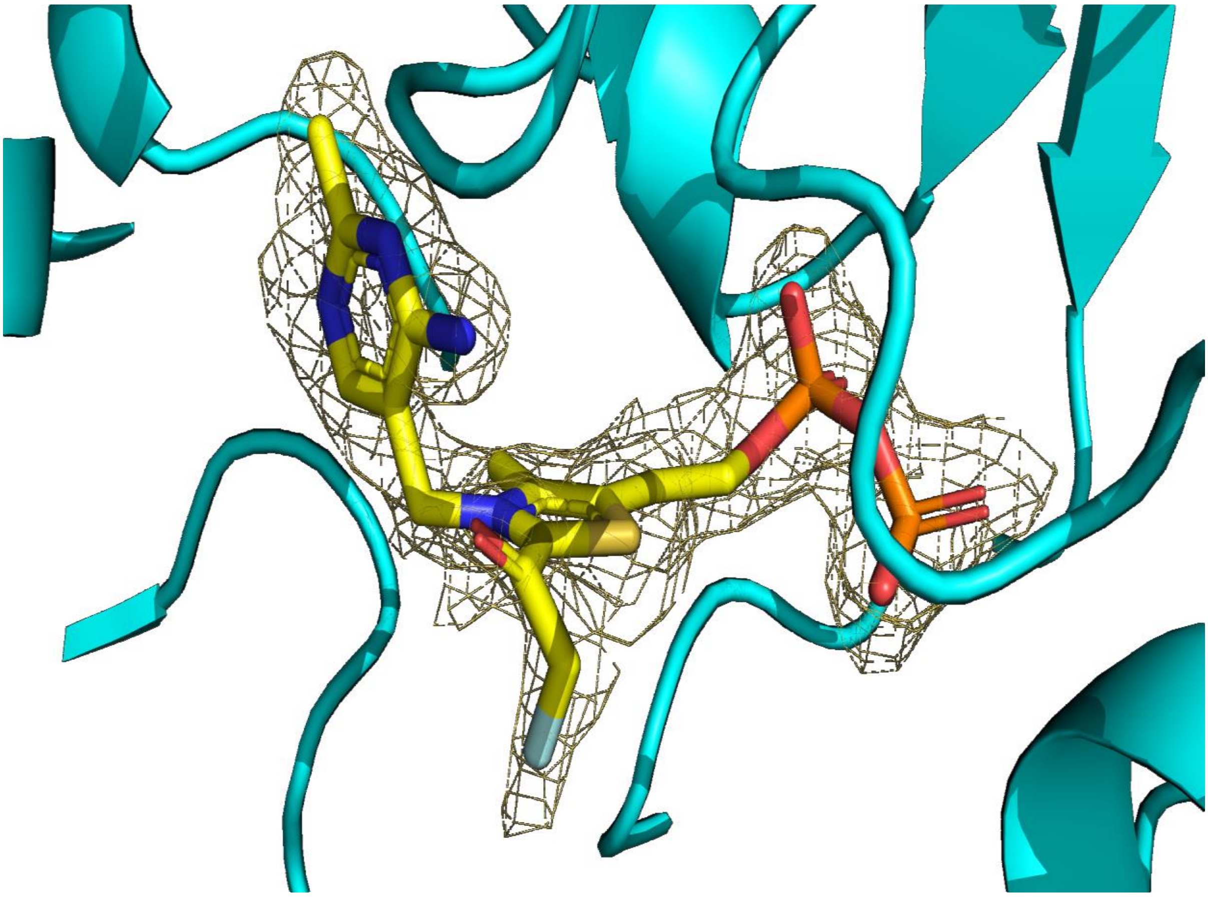

**Figure S4:** Composite omit electron density maps of 2-fluoroacetyl-ThDP binding to paDXPS, the difference electron density map of the ligand ($F_O$ - $F_C$) was contoured at 3$\sigma$ with phases calculated from a model that was refined in the absence of the ligand and is shown as a grey isomesh.

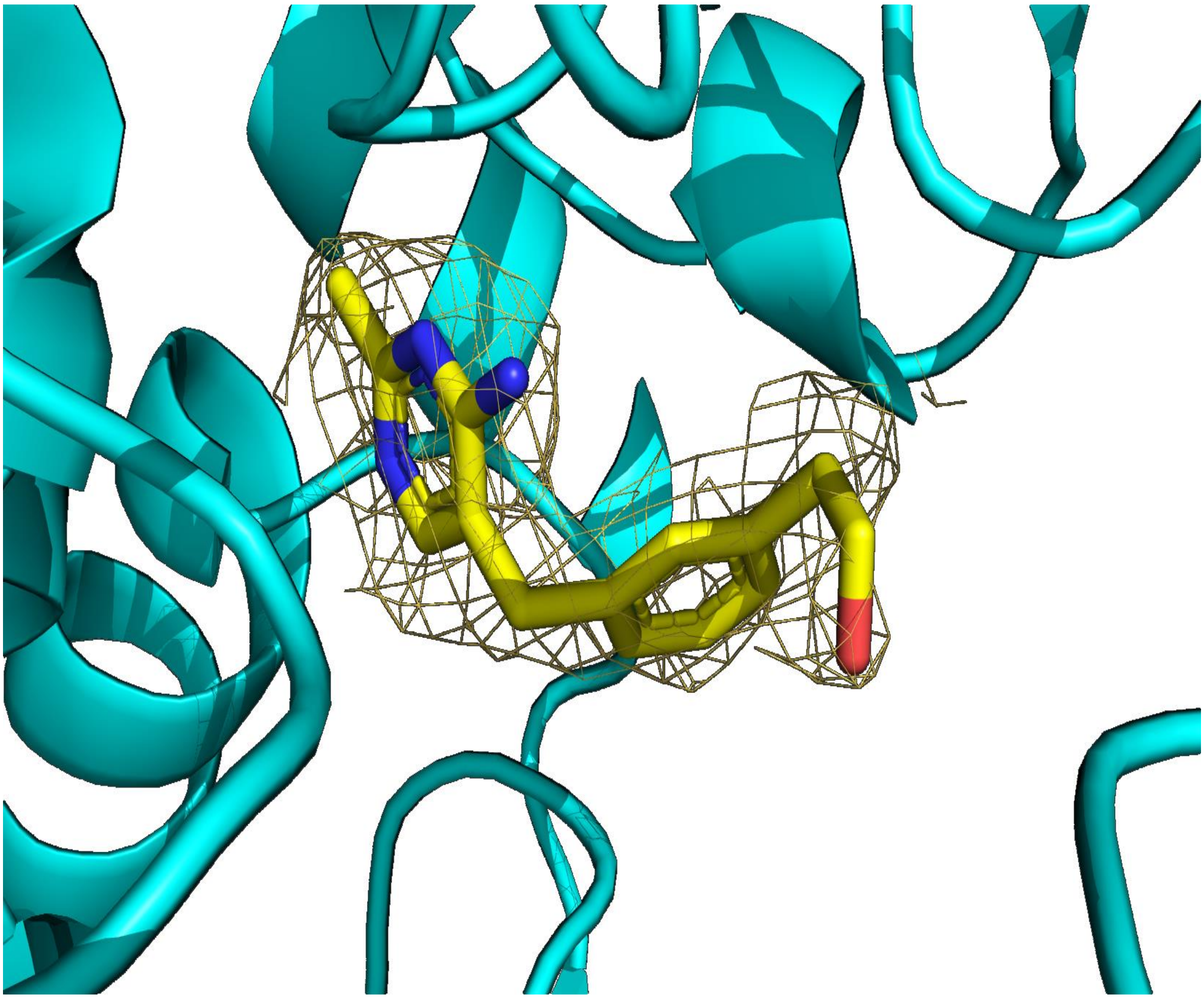

**Figure S5**: Composite omit electron density maps of inhibitor (1) to paDXPS, the difference electron density map of the ligand ($F_O$ - $F_C$) was contoured at 3$\sigma$ with phases calculated from a model that was refined in the absence of the ligand and is shown as a grey isomesh.
